# Immunological role of primary cilia of dendritic cells in human skin disease

**DOI:** 10.1101/2020.02.04.933333

**Authors:** Manami Toriyama, Defri Rizaldy, Motoki Nakamura, Fumitaka Fujita, Fumihiro Okada, Akimichi Morita, Ken J. Ishii

## Abstract

Primary cilia are a unique organelle, known to provide a signaling hub for variety of cell activities. Their potential role(s) in human immune homeostasis and diseases, however, have yet to be explored. Here, we show that human dendritic cells (DCs) express primary cilia-like structure. The primary cilia growth during DC proliferation by GM-CSF was shut off by DC maturation agents, suggesting the role of primary cilia to transduce proliferation signaling. PDGFRα pathway, one of proliferation signal in primary cilia, promoted DC proliferation in a dependent manner of intra-flagellar transport system. In epidermis with atopic dermatitis patients, aberrant ciliated langerhans cells and keratinocytes with showing immature state were observed that may play a potential role in inflammation and skin barrier disorder.

## Main

Primary cilia are unique organelles protruding into extracellular spaces from a basal body, and work as a platform for signaling pathways [1]. Intraflagellar transport (IFT) system is essential for axonome elongation and ciliary protein transport [2]. *IFT* gene mutations or disruption of this transport system eliminate primary cilia, resulting developmental defects, signaling defects and ciliopathy [3-5]. Primary cilia are formed when cells are in G0 or G1 phase, and their components regulate cell cycle progression [6]. Thus, primary cilium formation and the cell cycle tightly regulate each other; it is widely thought that primary cilia regulate cell proliferation and differentiation in many types of cells or tissues. While it has been well accepted that almost all types of cells can generate primary cilia, whether immune cells and/or stromal cells in immune organs/tissues express primary cilia and if so, what would be the precise roles its existence in the immune system and mechanisms of their action have not been explored until recently. In 2015, Prosser and Morrison reported that immortalized T and B cells have primary cilia [7]. Furthermore, Ezratty et al., reported that mouse embryonic epidermal keratinocytes (KC) has primary cilia, regulating KC differentiation [8].

Skin is the biggest tissue, which plays immunologically important roles in its homeostasis as well as pathological conditions such as injection and injury. Amongst three layers of the skin; epidermis, dermis, and subcutaneous tissue, KCs are the major cells in epidermis, and its proliferation and differentiation are strictly regulated to maintain skin homeostasis [9]. KCs finally form the stratum corneum working as physical barrier against pathogens [10]. Immune cells, including Langerhans cells (LCs), which have a similar role as dendritic cells (DCs), also exist in epidermis, and maintain skin homeostasis by working as antigen-presenting cells to activate T cells [11-13]. It has long been accepted that LCs have a pivotal role in bridging innate and acquired immunity, which is required for skin homeostasis. On the other hand, they are involved in the pathology of allergic skin diseases, including atopic dermatitis [14-16]. Yet, there are very few studies exploring the potential role of primary cilia in LC and/or other DCs of the skin in relation with the stromal cells such as KCs in either homeostasis or pathogenesis of skin immunity.

The prevalence of atopic dermatitis has increased greatly in the past 30 years, and it is widely known that environmental factors such as mite antigen in house dust can trigger atopic disease. In both the early and chronic stages of atopic dermatitis, type 2 immune responses, which are characterized as the elevation of several type 2 cytokines and IgE production, are dominant [17]. Chemokines secreted from KCs are upregulated, recruiting Th2 cells which strongly induce Th2 responses [18]. Atopic dermatitis often features skin barrier disruption, leading to dryness, itchiness, and invasion by pathogens such as *Staphylococcus aureus*. Topical steroids, tacrolimus, and moisturizers are use in its treatment, but because of its complicated pathogenesis, it often recurs after improvement. As such, it is clinically important to regulate inflammation and skin barrier maintenance, targeting both KCs and immune cells, however, potential roles of primary cilia in the pathogenesis of atopic dermatitis have not been fully clarified.

We therefore hypothesized that there may be potential role of primary cilia in the immune system in both homeostatic and pathological conditions, and examined whether human immune cells such dendritic cells express primary cilia, derived from primary blood monocytes and cell lines in vitro as well as human skin samples of healthy and atopic dermatitis patients. We further went on analyzing the molecular mechanism(s) by which expression and/or growth of primary cilia in human immune cells are regulated, and their physiological relevance in homeostasis and pathological condition of the skin.

## Results

### Human primary immune cells have primary cilium-like structures

To examine the primary cilia expression in human skin, we took healthy human skin samples and visualized the primary cilia in epidermis and dermis by staining with acetylated tubulin using fluorescent microscopy. As expected, we found many ciliated cells in dermis, which suggests that most ciliated cells were fibroblasts (extended data Fig. 1A). We found primary cilium-like structure in the epidermal basal area where proliferating KC stem cells are populous. Primary cilia-like structures were also detected in the stratum spinosum, at a lower frequency than in the basal layer (Fig. 1B). We hypothesized that most of the ciliated epidermal cells were KCs. However, terminal differentiation of KCs induces programmed cell death in the granular layer to form the stratum corneum [19]. As loss of cilia induces apoptosis in HeLa cells [20], we hypothesized that the ciliated cells in the striatum layer were not KCs but were resident living immune cells. When examining primary cilia in CD4+ T cells, CD8+ T cells, and Langerhans cells (LCs), we found only LCs in epidermis, positively merged with acetylated tubulin (Fig. 1B). Of note, most primary cilium-like structures in the stratum spinosum were not co-localized with langerin-positive cells. We did not determine the type of cells ciliated in this layer, but we hypothesized that they were KCs just before apoptosis. Mouse KCs have primary cilia ([8]), so human adult KCs may also be ciliated.

**Figure 1.**
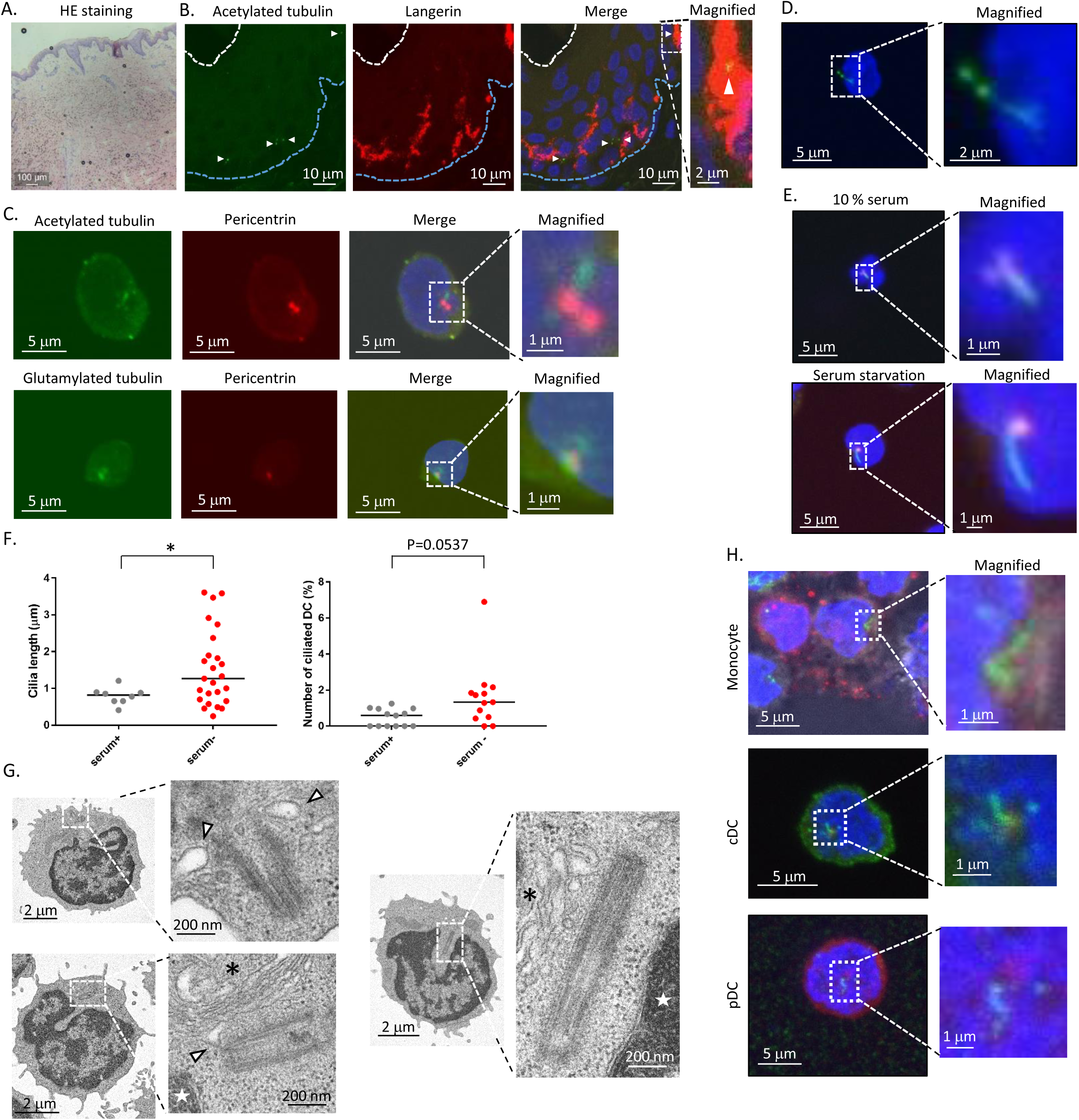
Human Primary immune cells have primary cilia-like structure. (A) HE staining of healthy human skin. (B) Langerin-positive langerhans cells and acetylated tubulin (primary cilia marker) in human healthy epidermis. Arrowhead indicates primary cilia-like structure. Blue dot line indicates epidermal basal layer. White dot line indicates stratum corneum. Dot box indicates magnified area shown in right. (C) Immunostaining image of human PBMC stained with acetylated tubulin or glutamylated tubulin. Dot box indicates magnified area shown in right. (D) Immunostaining image of human PBMC expressing Arl13B-GFP. Cells were electroporated with Arl13B-GFP expression plasmid, then immunostained by using anti GFP antibody. Dot box area was magnified. (E) Human PBMC was cultured in media supplemented with 10% FBS (serum +) or 0.5% FBS (Serum -) for 16 hrs, then immunostained with acetylated tubulin (Green), and pericentrin (Red). Dot box area were magnified. (F) Number of ciliated cells or cilia length shown in (E). Bar indicates median. *P<0.05. (Mann-Whitney U test). (G) Electron microscope image of primary cilia-like structure. Arrow head indicates ciliary vesicle like structure. Asterisk indicates goldi apparatus. Star indicates nucleus. Dot box indicates magnified area shown in right. (H) Monocyte, pDC and cDC were isolated from PBMC using flow cytometry, then isolated cells were immunostained with acetylated tubulin (green), and pericentrin (red).

LCs are a kind of DC and have a similar function as conventional DCs (cDCs). To see whether immune cells in blood, especially cDCs, are ciliated, we isolated peripheral blood mononuclear cells (PBMCs), a mixture of immune cells, from human peripheral blood, and immunostained them with acetylated or glutamylated tubulin. Nearly 2% of cells had primary cilium-like structures a showing single protrusion stained with stabilized tubulin, extending from the pericentrin by nearly 1 µm (Fig. 1C). We examined whether the structure was Arl13B positive. Arl13B is known as primary cilia marker, and is required for cilia formation and maintenance [21]. Arl13B-GFP was exogenously expressed in PBMCs, then cells were immunostained with green fluorescent protein (GFP). A GFP signal was observed as a single linear structure (Fig. 1D) similar to the stabilized tubulin (Fig. 1C). As the formation of primary cilia is strongly associated with the cell cycle and serum starvation promotes their elongation in many types of cells by inducing cell cycle arrest [22, 23], we examined whether human primary immune cells use a similar mechanism to elongate primary cilia. Serum starvation by culturing in 0.5% serum for 16 h significantly increased the frequency of primary cilium-like structures, which tended to be longer than those in cells treated in 10% serum (Fig. 1E, F). For further investigation of primary cilium-like structures in PBMCs in detail, we used transmission electron microscopy. We observed the vesicle–centrosome interaction, which resembles a ciliary vesicle (Fig. 1G). Also, centrosome elongation resembling axoneme extension, which is found in early primary cilium elongation, was clearly observed (Fig. 1G). These results raise the strong possibility that human immune cells are ciliated, and have similar machinery to elongate primary cilia.

To identify specific types of ciliated immune cells, we isolated monocytes, cDCs, plasmacytoid dendritic cells (pDCs), CD4 T cells, CD8 T cells, natural killer cells, and B cells from human PBMCs by flow cytometry and immunostained them with acetylated tubulin. Except for CD8 T cells, all other cells had primary cilium-like structures resembling that in Figure 1C (Fig. 1G, Supp. Table 1). cDCs had the highest rate, so we focused on the function of primary cilia in DCs. The frequency of ciliated primary DCs isolated from PBMCs was comparable to that of LCs in epidermis (Fig. 4D, Supp. Table 1). In summary, these results suggest the presence of primary cilia in DCs and LCs. We next sought the function of primary cilia in these cells.

**Figure 2.**
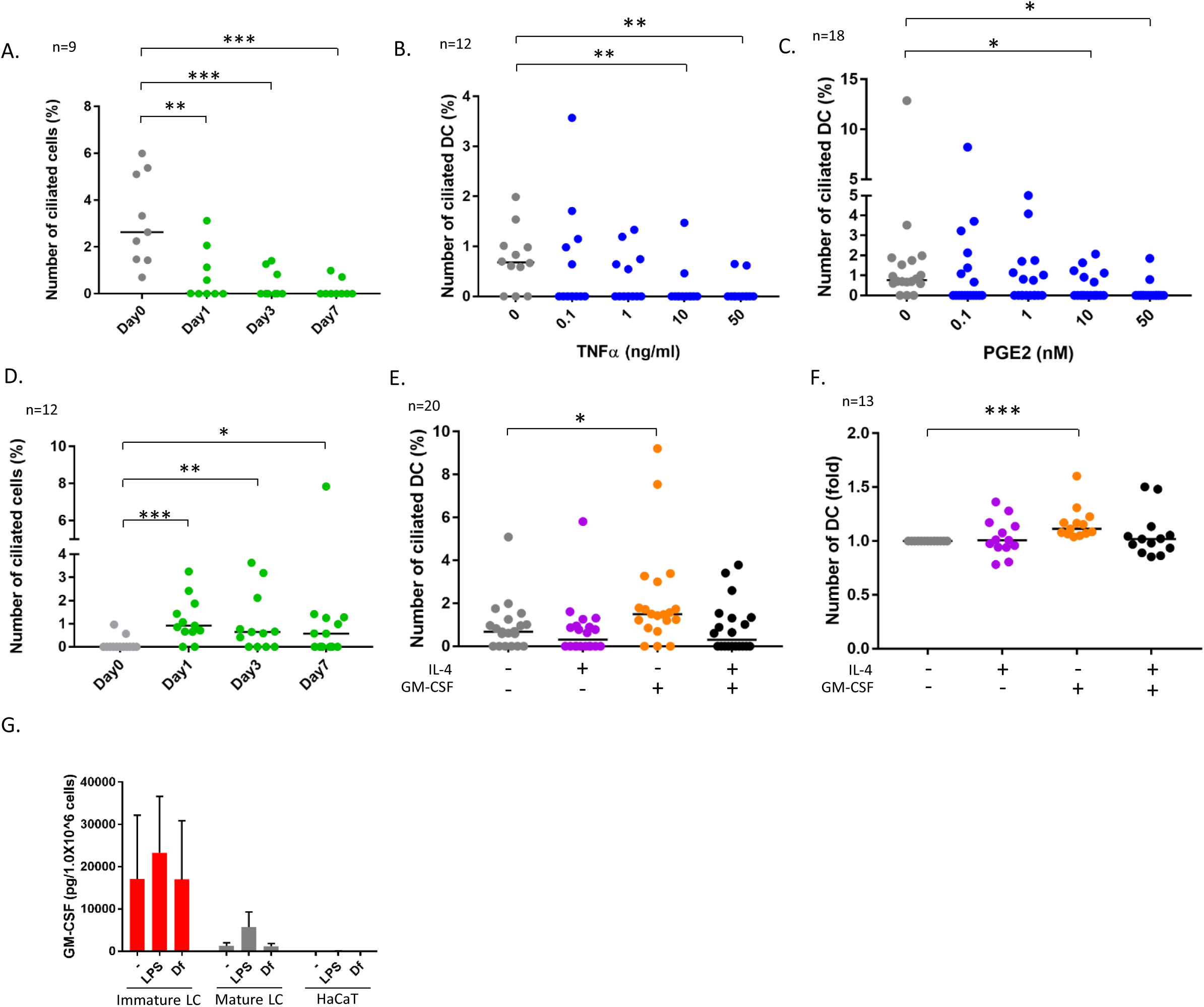
DC maturation decreases primary cilia. (A) CD14+ monocytes were isolated from human PBMC by magnetic beads, then differentiated into DC by stimulating with 50 ng/ml GM-CSF, 50 ng/ml IL-4 and 50 ng/ml TNFα. Percentage of ciliated cell was shown in graph. Bar indicates median. **P<0.01, ***P<0.001 (Mann-Whitney U test). n=9. (B, C) Human DC isolated from PBMC was stimulated with (B) TNFα, or (C) PGE2 for 24 hrs. Percentage of ciliated cells were shown in graph. Bar indicates median. *P<0.05, **P<0.01 (Mann-Whitney U test). n=12 and n=18, respectively. (D) CD14+ monocytes were isolated from human PBMC by magnetic beads, then differentiated into DC with 50 ng/ml GM-CSF and 50 ng/ml IL-4. Percentage of ciliated cell was shown in graph. Bar indicates median. *P<0.05, **P<0.01, ***P<0.001. (Mann-Whitney U test). n=12. (E) DCs isolated from PBMC were cultured for 24hrs with 50 ng/ml IL-4 and 50 ng/ml GM-CSF. Percentage of ciliated cells was counted and graphed. Bar indicates median. *P<0.05, (Fisher’s LSD). n=20. (F) DCs isolated from PBMC were cultured with 50 ng/ml IL-4, 50 ng/ml GM-CSF with CCK buffer for 48 hrs, then relative number of cells were calculated by measuring absorbance. Bar indicates median. ***P<0.001 (Mann-Whitney U test). n=13. (G) Average of GM-CSF expression measured by ELISA. Immature LC, Mature LC and HaCaT cells were stimulated with 10 μg/ml LPS or 10 μg/ml Df for 24 hrs in media supplemented with 0.5% serum. Culture supernatant was used for assay. Error bar shows SEM. n=5.

**Figure 3.**
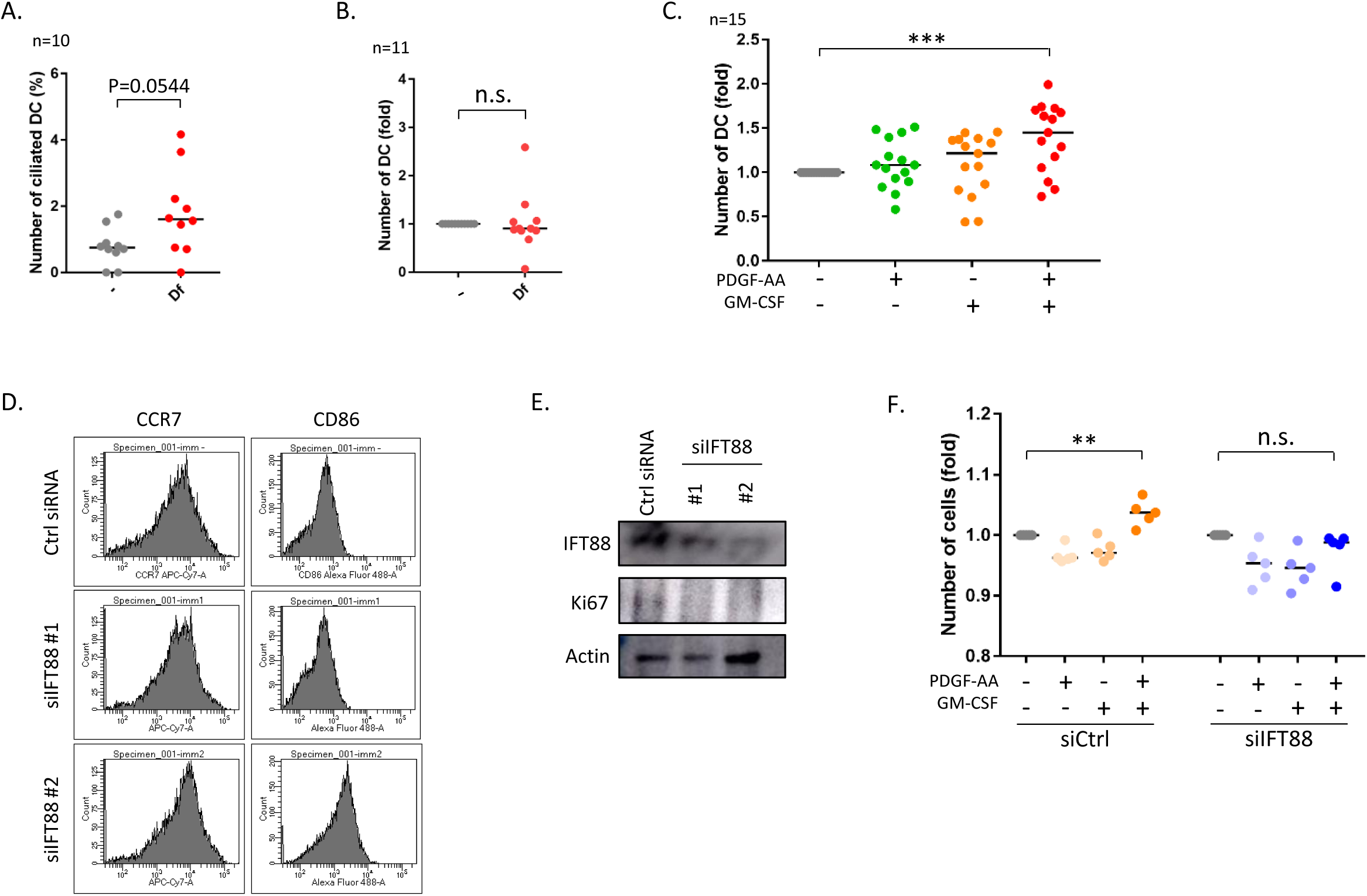
Primary cilia regulates proliferation and maturation via PDGFRα signaling. (A) DCs were cultured with 1 μg/ml Df for 48 hrs, then counted ciliated cell number. Bar indicates median. **P<0.01 (Mann-Whitney U test). (B) Relative number of DC was measured 48 hrs after stimulation with 1 μg/ml Df by measuring absorbance. Bar indicates median. *P<0.05 (Mann-Whitney U test). (C) Relative number of DC was measured 48 hrs after stimulation with 10 ng/ml PDGF-AA or 10 ng/ml GM-CSF. For the control of PDGF-AA stimulation, PDGF-AA solvent (4 mM HCl, 0.1% BSA) was added. Bar indicates median. Fisher’s LSD was performed. ***P<0.001 n=15. (D) To induce immature DC, THP1 cells were cultured with 100 ng/ml GM-CSF and 100 ng/ml IL-4 for 5 days. At day5, cells were electroporated with 100 nM siRNA targeting IFT88 or control siRNA. Two days after electroporation, expression of cell marker was measured by flow cytometry. (E) IFT88 and Ki67 expression in THP1-derived immature DC was investigated by western blotting. (F) Immature DC derived from THP1 was electroporated with 100 nM siRNA. Two days after electroporation, cells were stimulated with10 ng/ml PDGF-AA or 10 ng/ml GM-CSF for 48 hours. Proliferation activity was measured by measuring absorbance with adding CCK buffer. Cells were Bar indicates median. **P<0.01 (Student T test).

**Figure 4.**
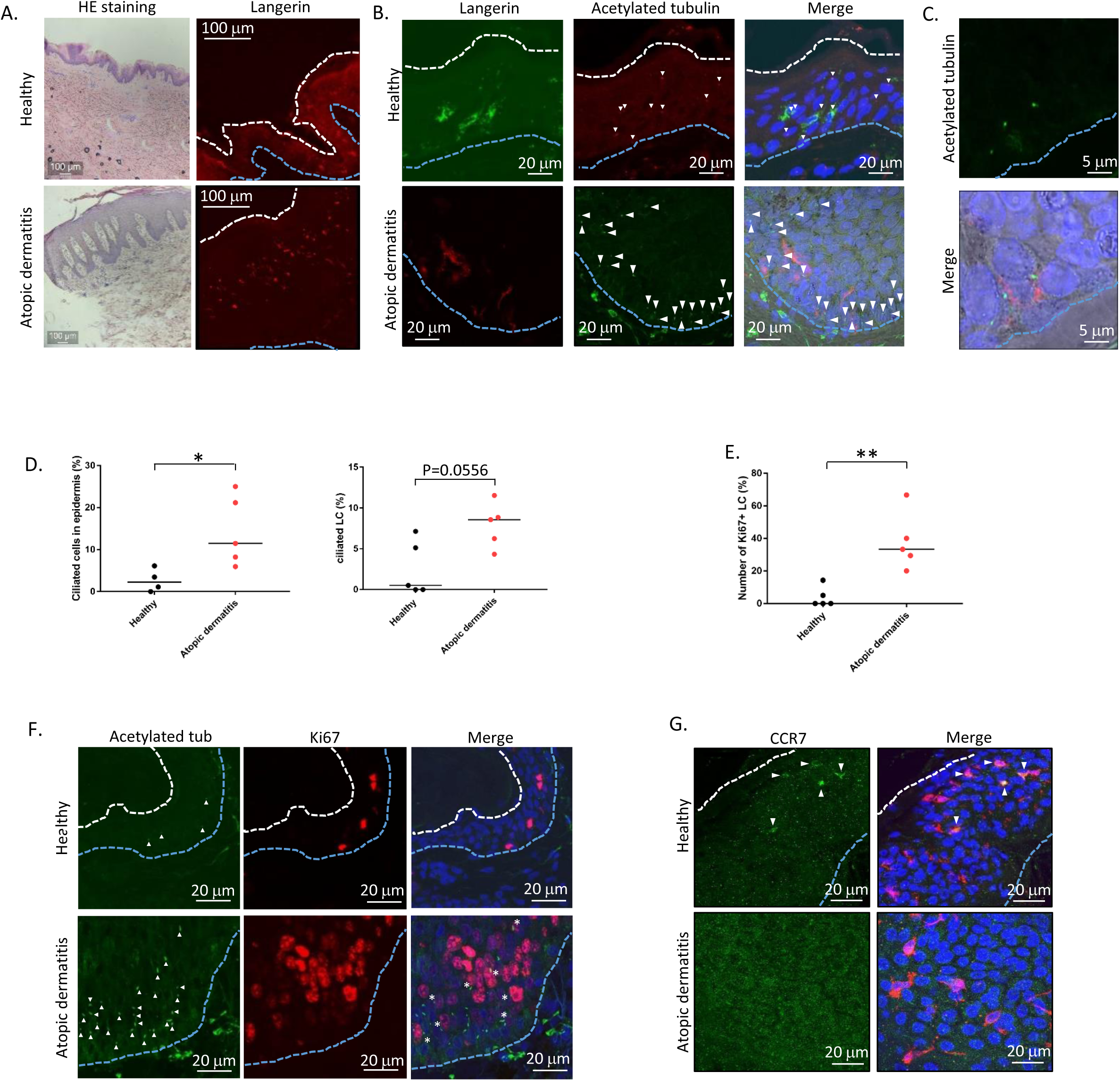
Primary cilia are increased in atopic dermatitis. (A) HE staining of human skin and Immunostaining of langerin in epidermis. Blue dot line shows basal layer. White dot line shows stratum corneum. (B) Langerin and acetylated tubulin staining in human epidermis. Blue dot line shows basal layer. Arrowhead indicates primary cilia-like structure. (C) Magnified image of ciliated LC in atopic skin. Langerin is shown in red. (D) Percentage of ciliated epidermal cells (upper), and ciliated LC (lower) in healthy or atopic dermatitis epidermis. Healthy samples; n=4 or n=5, atopic skin sample; n=5. Bar indicates median. *P<0.05 (Mann-Whitney U test). (E) ki67-positive LC in healthy and atopic skin were counted and graphed. *P<0.05 (Mann-Whitney U test). (F) Immunostaining for acetylated tubulin (green), and ki67 (red) in epidermis from healthy donor or atopic dermatitis patient. Blue dot line shows basal layer. White dot line shows stratum corneum. Arrow head shows primary cilia. Asterisk shows ki67-positive ciliated cells. (G) Immunostaining for CCR7 (green), and langerin (red) in epidermis. Arrow head indicates CCR7 positive LC

### DC maturation decreases primary cilia

DCs can be differentiated from monocytes *in vitro* [24]. To analyze whether monocyte-derived DCs are ciliated *in vitro*, we isolated CD14+ monocytes from human PBMCs and cultured them for 7 days with stimulation with GM-CSF, IL-4 and TNFα to differentiate into mature DCs. When we looked at primary cilia in them, the frequency of primary cilia was significantly decreased as day passed (Fig. 2A). To analyze whether DC maturation decreases primary cilium formation, we stimulated primary DCs with TNFα or PGE2 for 24 h. TNFα significantly decreased primary cilium formation dose-dependently (Fig. 2B). PGE2 stimulation had a similar effect (Fig. 2C), which suggests that DC maturation decreased primary cilium formation.

We also induced immature DCs by culturing with GM-CSF and IL-4. In immature DCs at day 7, nearly 1% of cells were ciliated, similar to the proportion in primary DCs (Fig. 2D, Supp. Table 1). The percentage did not differ significantly from day 1 to day 7 (Fig. 2D). Most DCs in healthy blood are immature, so these results suggest that immature DCs derived from CD14+ monocytes *in vitro* are similar to primary DCs, and raise the possibility that primary cilium formation was inhibited while monocytes differentiated into mature DCs.

Next, we investigated whether IL-4 or GM-CSF promoted primary cilium formation, because the frequency of primary cilia in immature DCs was a little higher than that in untreated (day 0) monocytes (Fig. 2D). GM-CSF significantly promoted primary cilium formation (Fig. 2E). Interestingly, IL-4 did not increase it, whereas IL-4 was simultaneously added with GM-CSF. It is widely known that cell maturation and cell proliferation show inverse behaviors. Immature or precursor cells proliferate much more than mature cells. To investigate the effect of GM-CSF and IL-4 on cell proliferation, we cultured immature DCs with GM-CSF or IL-4 and analyzed cell growth by MTT assay. GM-CSF increased DC proliferation, but IL-4 did not. Co-treatment with IL-4 canceled the effect of GM-CSF (Fig. 2F). These findings raise an interesting possibility that primary cilium formation is correlated with DC proliferation, making GM-CSF a candidate to promote primary cilium formation.

Having demonstrated that GM-CSF promoted primary cilium formation and cell proliferation, we evaluated which cells secrete GM-CSF. We stimulated CD14+ monocytes to differentiate into immature and mature LCs, then compared GM-CSF expression between those and human immortalized KCs (HaCaTs). Immature LCs expressed GM-CSF much more than mature LCs and HaCaTs did (Fig. 2G). Immature LCs spontaneously expressed GM-CSF without any stimulation, and stimulation did not alter its expression (Fig. 2G). IL-4 was not detected in LCs, but HaCaTs spontaneously expressed it (extended data Fig. 2). These results suggest that immature LCs in epidermis are the main producers of GM-CSF.

### Df promotes primary cilium formation

As GM-CSF promoted ciliogenesis and proliferation, we next focused on immunostimulants that promote GM-CSF secretion. GM-CSF is a Th2 cytokine, and its expression is elevated in atopic dermatitis [25], which can be triggered by house dust mite antigen. Therefore, we tested the effect of mite antigen on ciliogenesis. We also tested LPS, which activates TLR4 and induces strong Th1 immune responses in human immature DCs. We stimulated primary DCs with LPS, and mite antigen derived from *Dermatophagoides farinae* (*Df*). *Df* antigen slightly (but not significantly) increased the population of ciliated DCs (Fig. 3A). In contrast, LPS significantly decreased primary cilia (extended data Fig. 4A). We next analyzed how these immunostimulants stimulate cell proliferation activity. Contrary to expectation, LPS and *Df* treatment did not affect proliferation activity in DCs (Fig. 3B, extended data Fig. 3B).

KCs showed different responses to immunostimulants from DCs. LPS stimulation decreased KC proliferation while *Df* significantly increased it (extended data Fig. 3C). Ki67 expression in HaCaTs was increased as expected after stimulation with *Df* (extended data Fig. 3D). The differences in cell responses between primary DCs and HaCaTs are not surprising, but we hypothesized that Df is one of the important molecule to regulate DCs and LCs functions.

### PDGFRα signaling promotes DC proliferation

PDGFRα is highly localized in primary cilia [26]. PDGF-A, a specific ligand for PDGFRα, promotes fibroblast and KC proliferation during wound healing of skin [9]. We identified PDGFRα expression in primary DCs and HaCaTs (extended data Figs. 3D, 4B, 4D). To identify the type of cells secreting PDGF-A, we performed ELISA assay with culture supernatant of immature LCs, mature LCs, and HaCaTs. HaCaTs spontaneously expressed PDGF-A, but LCs did not. These results strongly suggest that KCs have a major role in secreting PDGF-A (extended data Fig. 4A).

Next we analyzed the function of PDGFRα signaling on proliferation in primary DCs and KCs. Co-treatment of DCs or KCs with PDGF-AA and GM-CSF significantly increased proliferation (Fig. 3C, extended data Fig. 4C, 4D). PDGFRα expression was not changed when cells were stimulated with PDGF-AA, GM-CSF, LPS and Df (extended data Fig. 3D, 4B, 4D). To investigate whether disruption of primary cilia causes DC maturation, we performed a knockdown experiment using siRNA targeting *IFT88*. Knockdown of *IFT88* increased the CD86 and CCR7 expression in THP1-derived immature DCs (Fig. 3D). Furthermore, ki67 expression was downregulated with *IFT88* knockdown (Fig. 3E). Interestingly, knockdown of *IFT88* canceled proliferation promoted by co-stimulation with PDGF-AA and GM-CSF (Fig. 3F). These results suggest an important role of the PDGFRα signaling pathway through primary cilia in DC proliferation.

### Primary cilia are increased in epidermis with atopic dermatitis

LCs proliferate extensively in epidermis with atopic dermatitis ([27], Fig. 4A). To address whether atopic condition alters primary cilium formation, we investigated the frequency of primary cilia in human epidermis with atopic dermatitis. Compared with healthy skin, the number of primary cilia was greatly increased especially in the basal area (Fig. 4B, D). Interestingly, ciliated cells other than LCs were often detected in atopic dermatitis (Fig. 4B,C). The frequency of ciliated cells (all types) and of ciliated LCs in atopic LCs was increased (Fig. 4D). We did not determine ciliated cell type other than LC, but we speculated they were KC because majority of epidermal cells are KC.

We also investigated the number of proliferating cells in atopic epidermis. Ki67 is a proliferation marker, and its expression is highly increased in S and M phases. In atopic dermatitis, ki67 positive LC is increased [27], and we found similar results. Ki67-positive epidermal cells were significantly increased in atopic epidermis (Fig. 4E). Unexpectedly, some ciliated cells were positively stained for ki67 in atopic epidermis, but none were positive in healthy skin samples (Fig. 4F). Primary cilia are generally formed in G0 or G1 phase, and their formation is repressed in proliferating cells in G2/M phase ([6]). We did not determine the specific cell cycle phase that cells were in, but strong expression of ki67 in atopic epidermis suggests that ki67 positive atopic epidermal cells were in S or G2/M phase. It is widely accepted that proliferation and maturation are highly correlated, so we next investigated an LC maturation marker, CCR7, in epidermis. In healthy epidermis, nearly 35% of LCs were positively stained with CCR7, but atopic LCs were not (Fig. 4G). This result strongly indicates that atopic LCs are immature.

We also explored the expression of KC maturation markers. In healthy epidermis, K14, a marker of immature KCs, was highly expressed near the basal layer, and K10, a marker of mature KCs, was highly expressed only in the stratum granulosum (Fig. 5A). In atopic epidermis, in contrast, K14 was highly expressed in the basal layer and moderately expressed throughout the stratum spinosum and stratum granulosum (Fig. 5B). Interestingly, K10 was strongly expressed even in the stratum spinosum, along with K14 (Fig. 5B). We next investigated other markers of differentiation, loricrin and filaggrin. These proteins are necessary for skin barrier formation, and both are decreased in more than 20% of patients. It is widely accepted that barrier disruption increases the risk of atopic dermatitis [28-30]. The percentage of ciliated cells was significantly increased in patients with low levels of loricrin, but was not correlated with filaggrin expression (Fig. 5C, extended data Fig. 5). Interestingly, examining the correlation between IgE level and Loricrin expression, as well IgE level and primary cilia did not have significant differences (Fig. 5D, E). These results suggest the physiological importance of primary cilia in KCs to maintain adequate loricrin expression and skin barrier formation, which develops but not exacerbate atopic dermatitis. In summary, immunostaining results strongly suggested excessive proliferation and abnormal primary cilium formation in atopic dermatitis, which in turn caused hyperproliferation and sustenance of immature LCs and KCs in atopic dermatitis.

**Figure 5.**
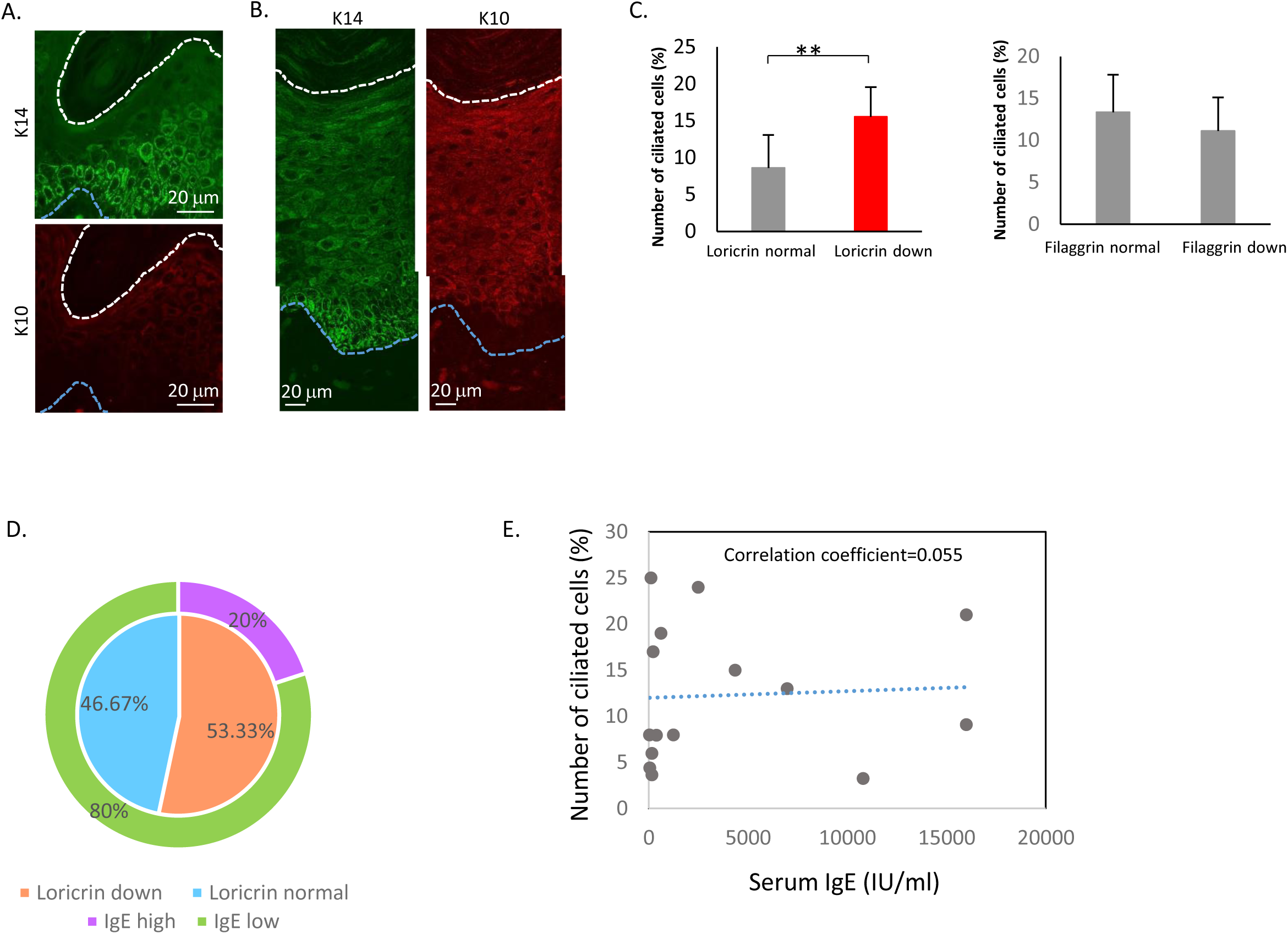
Keratinocyte differentiation marker, Loricrin expression correlates with primary cilia rate. (A,B) Immunostaining for K14 (green) and K10 (red) in (A) healthy epidermis, and in (B) atopic epidermis. (C) Correlation between primary cilia and epidermal barrier Protein, loricrin and Filaggrin. Atopic dermatitis epidermis were immunostained with Loricrin or Filaggrin, with acetylated tubulin, then calculated percentage of ciliated cells. (D) Percentage of atopic dermatitis patients showing Loricrin normal, loricrin down, IgE low, IgE high, respectively. (E) Correlation between primary cilia percentage and serum IgE level in atopic dermatitis patients.

### Immature Langerhans cells secrete chemokines

Our findings show that ciliated DCs and LCs were immature and highly proliferative *in vitro* and in atopic dermatitis. We next sought the physiological function of immature LCs. Th2 T cells highly infiltrate atopic epidermis and secrete Th2 cytokines [18]. Thus, we asked whether immature LCs secrete chemokines to recruit Th2 T cells. To answer this, we analyzed chemokine expression. MCP1 was highly secreted by immature LCs relative to mature LCs and KCs (extended data Fig. 6A). In contrast, TARC/CCR17 was expressed by mature LCs, but not so much by immature LCs or KCs (extended data Fig. 6B). MDC was highly expressed in both immature and mature LCs but was not detected in KCs (extended data Fig. 6C). These data suggest the important role of LCs but not KCs in secreting chemokines to recruit Th2 cells. In atopic dermatitis, immature LCs were highly increased (Fig. 4G). Our results raise the strong possibility that immature LCs are the main releasers of Th2 chemokines in atopic dermatitis, which exacerbate disease.

## Discussion

We identified primary cilia in human epidermis and immune cells from blood (Fig. 1). Primary cilium formation in epidermis was highly promoted in atopic dermatitis (Fig. 4). High correlation of primary cilium formation and proliferation suggests a primary function of cilia in epidermal cell proliferation in atopic dermatitis. In atopic epidermis, ki67-positive ciliated cells were greatly increased. Primary cilium formation is basically inhibited in G2/M and S phase, and this phenotype was not found in healthy epidermis, so the cell cycle and proliferation may be disrupted in atopic dermatitis which maybe a cause of disease. Our data suggested that GM-CSF promotes primary cilium formation, and PDGFRα signaling via primary cilia promotes proliferation of DCs (Figs. 2-4). The candidate of atopic dermatitis inducer, Df, induced primary cilia formation in DCs and proliferation in KCs. Because KCs produce PDGF-AA constantly, Df may induce thickening of epidermis, which strongly induces LC proliferation. We have not identified what induces atopic dermatitis, however, we propose that GM-CSF and Df are strong candidates to develop atopic dermatitis by regulating primary cilia formation and proliferation. Interestingly, ciliopathy patients in several pedigrees show frequent atopic dermatitis or asthma along with other phenotypes caused by primary cilium defect [31, 32]. We have not identified genetic alteration of cilium-related genes in atopic dermatitis patients, so further experiments will be required.

Interestingly, the purchased HaCaTs and primary KCs were not ciliated *in vitro*, although epidermal KCs in tissue were (data not shown; Figs. 1, 4). Because of this technical difficulty, we could not study the function of primary cilia in KCs in detail. The differences between epidermal KCs and isolated KCs are unknown, but 3D cell-cell interaction or environmental factors in tissue may be involved in primary cilium formation. Further investigation is required to explain the function of primary cilia in KCs.

Proliferation and maturation are highly inversely correlated with each other. Therefore, we asked whether inhibition of proliferation by disrupting primary cilia promoted maturation. We demonstrated that knockdown of *IFT88* in THP1-derived DCs promoted maturation by attenuating proliferation activity promoted by PDGFRα signaling (Fig. 3D, E). These results raise a strong possibility that PDGFRα signaling in primary cilia regulates cell proliferation and inhibits maturation. We tried to identify the expression of PDGFRα in both healthy and atopic epidermis, but the experiment did not work (data not shown). Furthermore, we could not detect the PDGFRα localization in primary cilia in LCs and DCs, even though these cells expressed PDGFRα detected by western blotting (extended data Figs. 4, 5). Interestingly, we found constitutive expression of PDGF-AA in HaCaTs (extended data Fig. 5A). This result raises a possibility that PDGF-AA expression is higher in atopic skin because the number of KCs is obviously increased. The relationship between pathophysiology and PDGF-AA expression will be the next topic of research. Imatinib, a PDGFRα antagonist, treats asthma by decreasing mast cell activation in patients and by decreasing MCP1 expression in mice [33, 34]. Its effectiveness in the treatment of atopic dermatitis has not been elucidated, but these reports suggest the importance of PDGFRα signaling in allergic disorders. Furthermore, methotrexate, a folate antagonist, successfully treated atopic dermatitis [35, 36]. Previously we demonstrated that folate metabolic pathway is required for primary cilium formation [37]. These knowledge support our idea that the regulation of primary cilia or PDGFRα signaling could help in the treatment of atopic dermatitis.

On the basis of our novel findings, we propose that primary cilia in LCs or in KCs have an important role in skin homeostasis by regulating proliferation, and excess cilium formation may cause atopic dermatitis. Over-formation of primary cilia in KCs sustains immaturity, impairing barrier function by reducing loricrin expression. Ciliated immature LCs strongly recognize antigens or immunostimulants that pass through the skin barrier, and LCs secrete GM-CSF. GM-CSF promotes primary cilium formation in both KCs and LCs, and proliferation signals, including PDGFRα, are transduced more strongly. Chemokines secreted from immature LCs strongly recruit Th2 T cells, which causes a vicious cycle of atopic dermatitis (extended data Fig. 6D).

## Methods

### Human skin samples

#### Isolation of primary DCs and CD14+ monocytes

Samples of human whole peripheral blood were purchased from the Japanese Red Cross Society according to the *Guidelines on the Use of Donated Blood in R&D, etc.* Blood from 89 healthy donors was drawn into a heparinized syringe and diluted with an equal volume of phosphate-buffered saline (PBS). A 35-mL volume of diluted blood was layered over 15 mL of Ficoll-Paque Plus density gradient medium (GE Healthcare) in Leucosep 50-mL tubes (Greiner Bio-One), then centrifuged at 500× *g* for 20 min at room temperature without braking. The PBMC fraction was carefully collected by pipetting, and the cells were washed with RPMI1640 supplemented with 10% fetal bovine serum (FBS). After centrifugation at 500× *g* for 10 min at room temperature, red blood cells were lysed with Ammonium-Chloride-Potassium (ACK) buffer (150 mM NH_4_Cl 10 mM KHCO3, 0.1 mM EDTA in PBS) in a conical tube at room temperature for 10 min. After washing with RPMI1640 containing 10% FBS, CD14-positive monocytes were isolated by CD14 microbeads, human (130-050-201, Miltenyi Biotec). The positive fraction was used as CD14-positive monocytes. DCs were isolated with a Blood Dendritic Cell Isolation Kit II, human (130-091-379, Miltenyi Biotec), from the negative fraction of the CD14-positive fraction.

#### Cell culture

HaCaT cells were cultured in Dulbecco’s modified Eagle’s medium (DMEM) supplemented with 10% FBS, 1% penicillin and 1% streptomycin. Immature and mature LCs were induced to differentiate from CD14-positive monocytes derived from human primary PBMCs. To induce immature LCs, CD14-positive monocytes were cultured in RPMI1640 supplemented with 10 ng/mL IL-4 (100-09, Shenandoah), 100 ng/mL GM-CSF (100-08, Shenandoah), and 10 ng/mL TGFβ (240-B, R&D Systems). To induce mature LCs, CD14-positive monocytes were cultured in RPMI1640 supplemented with 10 ng/mL IL-4, 100 ng/mL GM-CSF, 10 ng/mL TGFβ, and 20 ng/mL TNFα (MAN0003622, Gibco). A half volume of fresh medium was added every 2 days.

#### Isolation of monocytes, pDCs, and cDCs by flow cytometry

Human PBMCs were labeled with Alexa 647 anti-human CD11c (301620, clone 3.9, BioLegend), anti-human HLA-DR (Class III) PE-Texas conjugate (MHLDR17, Life Technology), BV421 anti-human CD14 (325627, clone HCD14, BioLegend), PE-Cy7 anti-human CD123 (560826, clone 7G3, BD Biosciences), APC-Fire 750 anti-human CD8a (301065, clone RPA-T8, BioLegend), APC-Fire 750 anti-human CD20 (302357, clone 2H7, BioLegend), and APC-H7 anti-human CD3 (560275, clone SK7, BD Pharmingen) in a Live/Dead Fixable Aqua Dead Cell Stain Kit (L34966, Invitrogen). Labeled cells were analyzed and isolated using BD FACSAria II (BD Biosciences). HLA-DR^high^–CD14^low^ cells were gated from a CD3-, CD8-, CD20-negative live-cell population and identified as the DC population. CD123^middle^–CD11c^high^ cells in the DC population were determined as cDCs. CD123^high^–CD11c^low^ cells in the DC population were determined as pDCs. CD14^high^–HLA-DR^high^–CD11c^high^–CD123^middle^ cells were determined as monocytes.

#### ELISA

In 48-well plates, 2.0 × 10^5^ HaCaT cells were stimulated with 100 µL of DMEM supplemented with 0.5% FBS. In 96-well plates, 2.0 × 10^4^ LCs were stimulated with 100 mL of RPMI1640 supplemented with 0.5% FBS. After 24 h stimulation, 0.5% Triton X-100 was added into the cell cultures and then cell lysate was collected with supernatant. Cell lysate was used for ELISA assay following the kit protocol. ELISA kits (CCL17, DY364, MCP1, DCP00, MDC, DMD00, IL-4, D4050, GM-CSF, DY215) were purchased from R&D Systems. PDGF-AA ELISA kits (EHPDGF) were purchased from Thermo Fisher Scientific. Absorbance (excitation, 450 nm) was measured on a microplate reader (Infinite F200 Pro, Tecan). As reference, background absorbance was measured at 560 nm.

#### Immunostaining

Paraffin-embedded human skin tissue was deparaffinized in xylene 2 times for 10 min each. The samples were then immersed in 100% ethanol, 95% ethanol, and 2 lots of deionized water for 10 min each. After rehydration, antigens were retrieved: The samples were immersed in 1 mM EDTA in deionized water and boiled by microwave for 15 min. They were then incubated with blocking buffer (10% FBS, 0.1% Triton X-100 in PBS) at room temperature for 1 h. First antibodies (listed below) were diluted 1:1000 in PBS for the detection of acetylated tubulin, langerin, and ki67. CCR7 antibody was diluted 1:100 and incubated at 4 °C overnight. After 3 washes in wash buffer (0.1% Tween-20 in PBS), second antibodies (Alexa 488-conjugated IgG, Alexa 594-conjugated IgG) containing 1/5000 Hoechst 33342 (Thermo Fisher Scientific) were reacted at room temperature for 2 h. After rinsing in wash buffer, samples were mounted with ProLong Gold antifade reagent (Thermo Fisher Scientific). For single-cell immunostaining following the blocking step, cells were mounted on MAS-coated slide glass (Matsunami). First antibodies comprised anti-acetylated α-tubulin (T7451, clone 6-11B-1, Sigma), anti-langerin (ab192027, clone EPR15863, Abcam), anti-ki67 (#9449, clone 8D5, Cell Signaling Technology; ab16667, clone SP6, Abcam), anti-CCR7 (MAB197, clone 150503, R&D Systems), anti-pericentrin (A301-348A, Bethyl), and anti-GFP (SC-9996, clone B-2, Santa Cruz Biotechnology) antibodies.

The number of ciliated cells was counted and statistically analyzed in Prism 7 software.

#### Proliferation assay

In 96-well plates, 3000 HaCaT cells or 5000 primary DCs were stimulated with reagent in 100 µL of medium containing 0.5% FBS and 10 µL of CCK-8 buffer (Cell Counting Kit-8, Dojindo). After 24 h, absorbance at 450 nm was measured on a microplate reader (Infinite F200 Pro). As reference, background absorbance was measured at 560 nm.

#### Electron microscopy

Cells were fixed with 2% glutaraldehyde in 0.1 M phosphate buffer (PB), pH 7.4, at 4 °C overnight. The fixed samples were washed 3 times with 0.1 M PB for 30 min each, and were post-fixed with 2% OsO_4_ in 0.1 M PB at 4 °C for 1 h. They were then dehydrated in a graded ethanol solution at 50% and 70% for 5 min each at 4 °C, 90% for 5 min at room temperature, and 3 changes of 100% for 5 min each at room temperature. The samples were infiltrated with propylene oxide (PO) 2 times for 5 min each, put into a 70:30 mixture of PO and resin (Quetol-812; Nisshin EM Co.) for 10 min, and then exposed to the open air overnight for the PO to volatilize. The samples were transferred to fresh 100% resin and polymerized at 60 °C for 48 h. The polymerized resins were ultrathin-sectioned at 70 nm with a diamond knife on an ultramicrotome (Ultracut UCT; Leica), and the sections were mounted on copper grids. They were stained with 2% uranyl acetate at room temperature for 15 min, washed with distilled water, and then secondary-stained with lead stain solution (Sigma-Aldrich) at room temperature for 3 min. The grids were observed by transmission electron microscope (JEM-1400 Plus; JEOL Ltd.) at an acceleration voltage of 100 kV. Digital images (3296 × 2472 pixels) were taken with a CCD camera (EM-14830RUBY2; JEOL Ltd.).

Reference mendeley

## Acknowledgments

The authors thank the Japanese Red Cross Society for supplying blood samples under the *Guidelines on the Use of Donated Blood in R&D, etc.* This work was supported in part by JSPS KAKENHI grant JP 19K17797 (to M.T.). The authors declare no conflicts of interest.

## Author contributions

M.T. and K.J.I. conceived and designed the experiments and wrote the manuscript; M.T., D.R., and M.N. conducted the experiments; and F.F., F.O., and A. Morita contributed to the writing and editing of the manuscript.

