## Supplemental figs and table for "Immunological role of primary cilia of dendritic cells in human skin disease"

### Slide 1
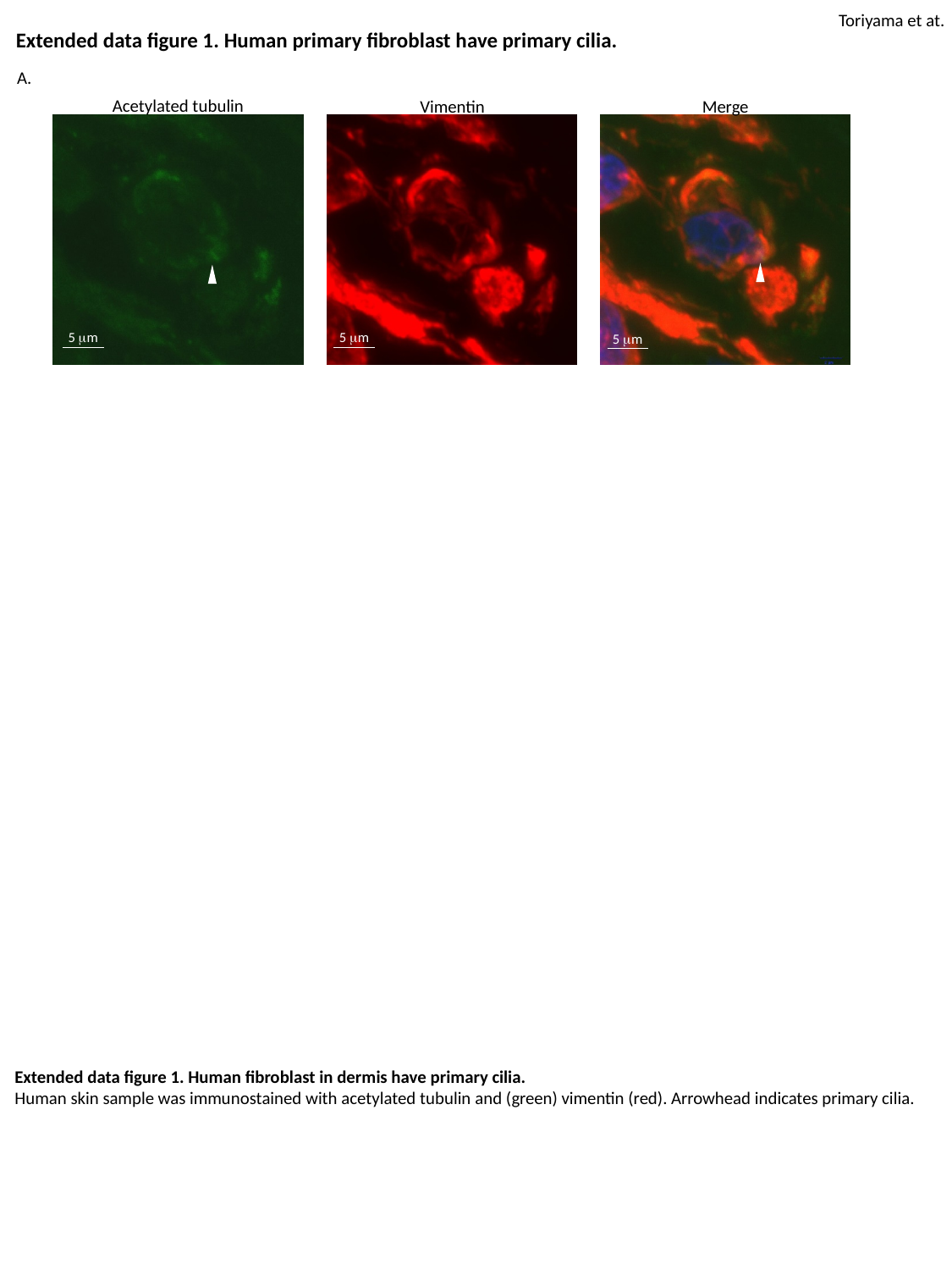

Toriyama et at.
Extended data figure 1. Human primary fibroblast have primary cilia.
A.
Acetylated tubulin
Merge
Vimentin
5 μm
5 μm
5 μm
Extended data figure 1. Human fibroblast in dermis have primary cilia.
Human skin sample was immunostained with acetylated tubulin and (green) vimentin (red). Arrowhead indicates primary cilia.

### Slide 2
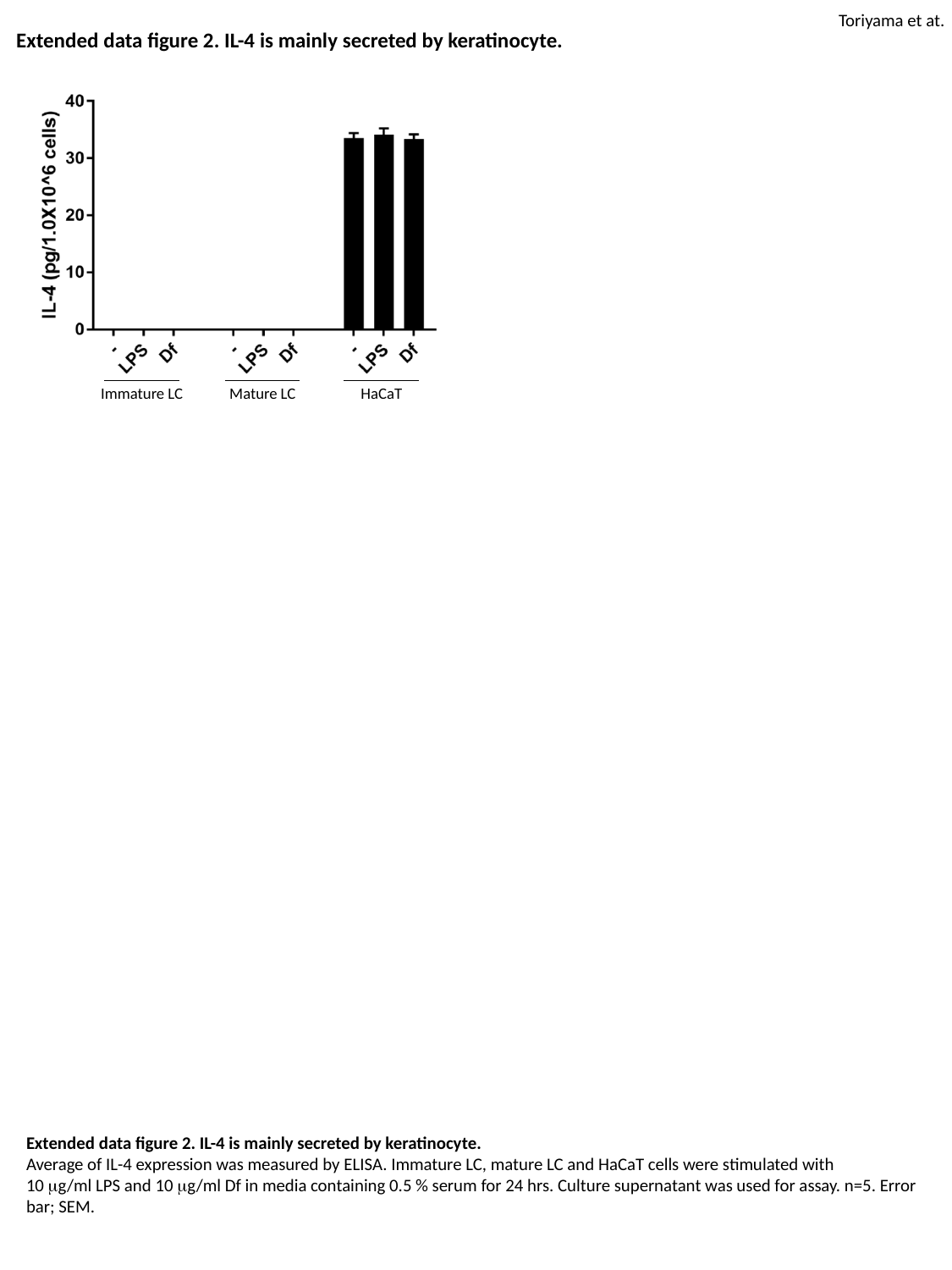

Toriyama et at.
Extended data figure 2. IL-4 is mainly secreted by keratinocyte.
Immature LC
Mature LC
HaCaT
Extended data figure 2. IL-4 is mainly secreted by keratinocyte.
Average of IL-4 expression was measured by ELISA. Immature LC, mature LC and HaCaT cells were stimulated with
10 μg/ml LPS and 10 μg/ml Df in media containing 0.5 % serum for 24 hrs. Culture supernatant was used for assay. n=5. Error bar; SEM.

### Slide 3
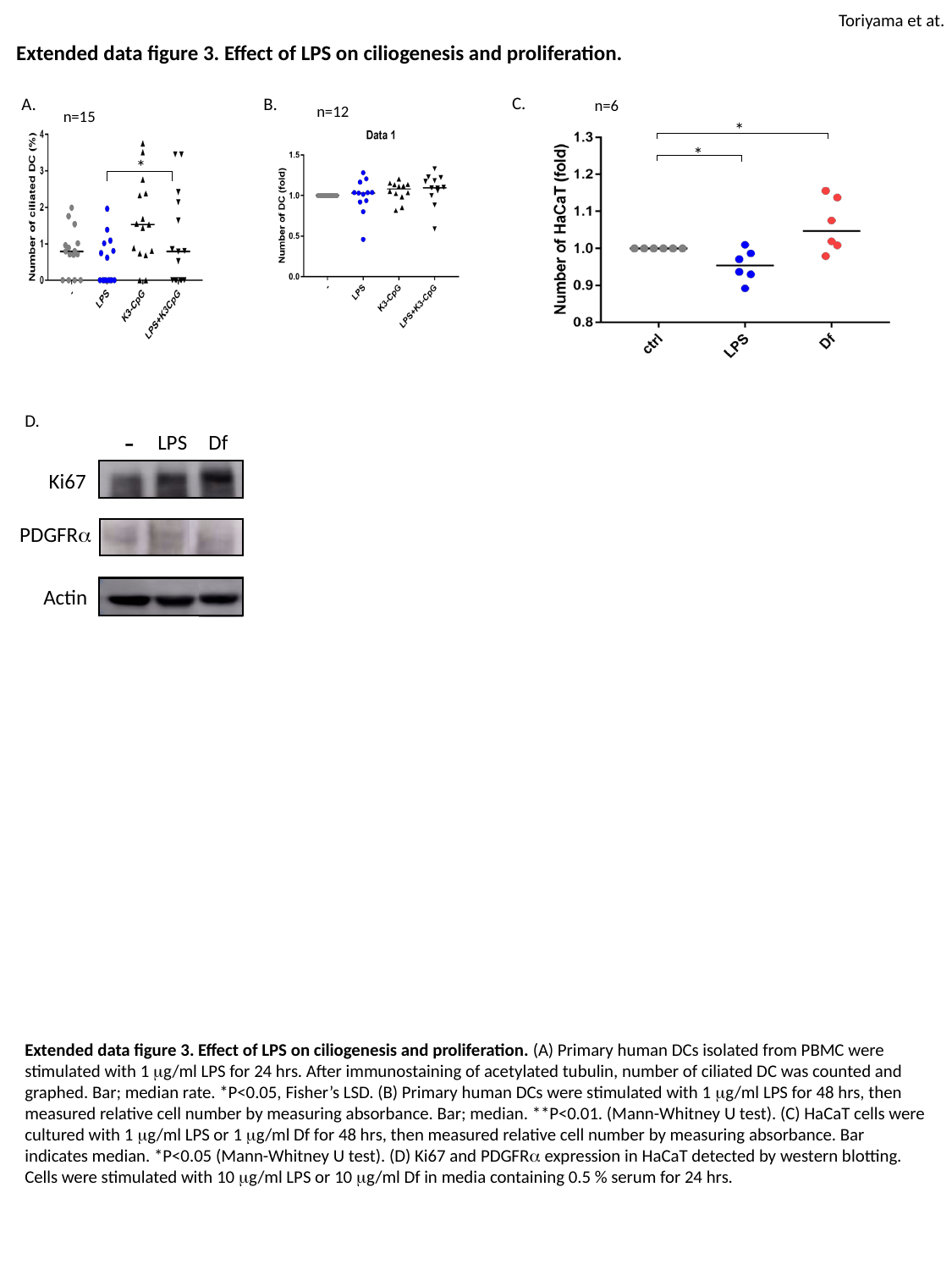

Toriyama et at.
Extended data figure 3. Effect of LPS on ciliogenesis and proliferation.
C.
B.
A.
n=6
n=12
n=15
*
*
*
D.
-
LPS
Df
Ki67
PDGFRα
Actin
Extended data figure 3. Effect of LPS on ciliogenesis and proliferation. (A) Primary human DCs isolated from PBMC were stimulated with 1 μg/ml LPS for 24 hrs. After immunostaining of acetylated tubulin, number of ciliated DC was counted and graphed. Bar; median rate. *P<0.05, Fisher’s LSD. (B) Primary human DCs were stimulated with 1 μg/ml LPS for 48 hrs, then
measured relative cell number by measuring absorbance. Bar; median. **P<0.01. (Mann-Whitney U test). (C) HaCaT cells were cultured with 1 μg/ml LPS or 1 μg/ml Df for 48 hrs, then measured relative cell number by measuring absorbance. Bar indicates median. *P<0.05 (Mann-Whitney U test). (D) Ki67 and PDGFRα expression in HaCaT detected by western blotting. Cells were stimulated with 10 μg/ml LPS or 10 μg/ml Df in media containing 0.5 % serum for 24 hrs.

### Slide 4
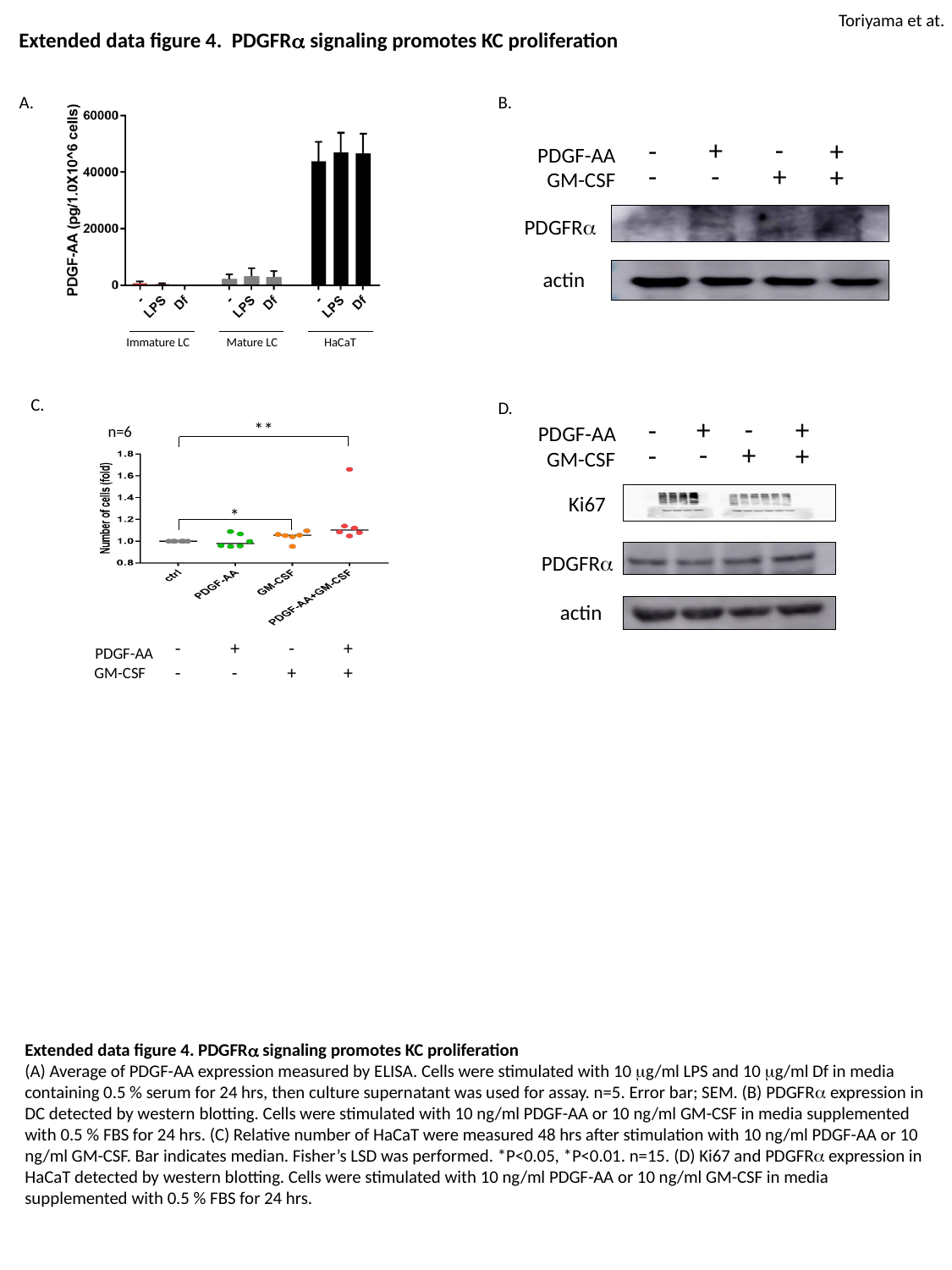

Toriyama et at.
Extended data figure 4. PDGFRα signaling promotes KC proliferation
A.
B.
+
-
+
-
PDGF-AA
-
+
-
+
GM-CSF
PDGFRα
actin
Immature LC
Mature LC
HaCaT
C.
D.
+
-
+
-
**
PDGF-AA
n=6
-
+
-
+
GM-CSF
Ki67
*
PDGFRα
actin
-
+
-
+
PDGF-AA
-
-
+
+
GM-CSF
Extended data figure 4. PDGFRα signaling promotes KC proliferation
(A) Average of PDGF-AA expression measured by ELISA. Cells were stimulated with 10 μg/ml LPS and 10 μg/ml Df in media containing 0.5 % serum for 24 hrs, then culture supernatant was used for assay. n=5. Error bar; SEM. (B) PDGFRα expression in DC detected by western blotting. Cells were stimulated with 10 ng/ml PDGF-AA or 10 ng/ml GM-CSF in media supplemented with 0.5 % FBS for 24 hrs. (C) Relative number of HaCaT were measured 48 hrs after stimulation with 10 ng/ml PDGF-AA or 10 ng/ml GM-CSF. Bar indicates median. Fisher’s LSD was performed. *P<0.05, *P<0.01. n=15. (D) Ki67 and PDGFRα expression in HaCaT detected by western blotting. Cells were stimulated with 10 ng/ml PDGF-AA or 10 ng/ml GM-CSF in media supplemented with 0.5 % FBS for 24 hrs.

### Slide 5
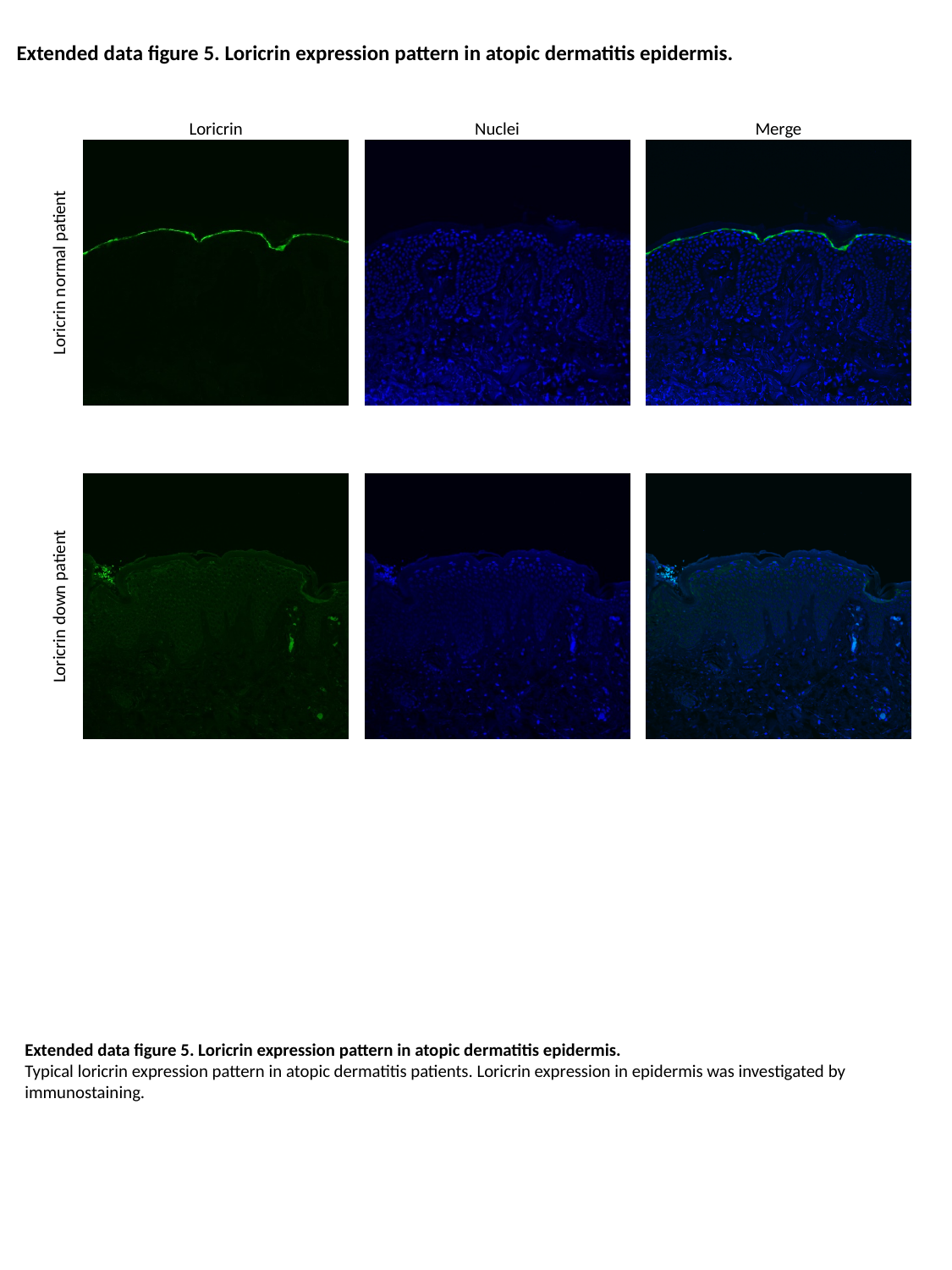

Extended data figure 5. Loricrin expression pattern in atopic dermatitis epidermis.
Loricrin
Nuclei
Merge
Loricrin normal patient
Loricrin down patient
Extended data figure 5. Loricrin expression pattern in atopic dermatitis epidermis.
Typical loricrin expression pattern in atopic dermatitis patients. Loricrin expression in epidermis was investigated by immunostaining.

### Slide 6
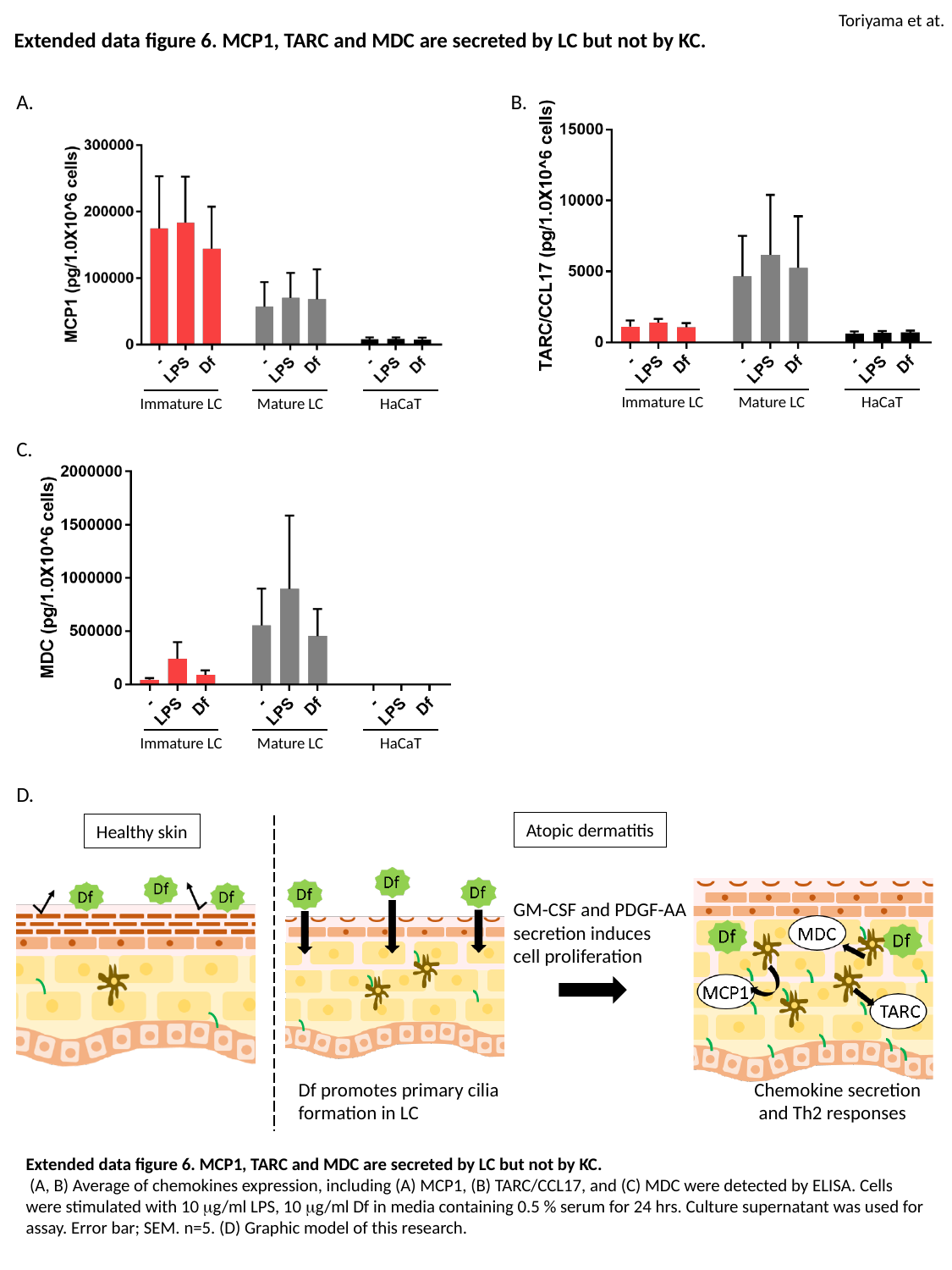

Toriyama et at.
Extended data figure 6. MCP1, TARC and MDC are secreted by LC but not by KC.
A.
B.
Immature LC
Mature LC
HaCaT
Immature LC
Mature LC
HaCaT
C.
Immature LC
Mature LC
HaCaT
D.
Atopic dermatitis
Healthy skin
GM-CSF and PDGF-AA
secretion induces
cell proliferation
Df promotes primary cilia
formation in LC
Chemokine secretion
 and Th2 responses
Extended data figure 6. MCP1, TARC and MDC are secreted by LC but not by KC.
 (A, B) Average of chemokines expression, including (A) MCP1, (B) TARC/CCL17, and (C) MDC were detected by ELISA. Cells were stimulated with 10 μg/ml LPS, 10 μg/ml Df in media containing 0.5 % serum for 24 hrs. Culture supernatant was used for assay. Error bar; SEM. n=5. (D) Graphic model of this research.

### Slide 7
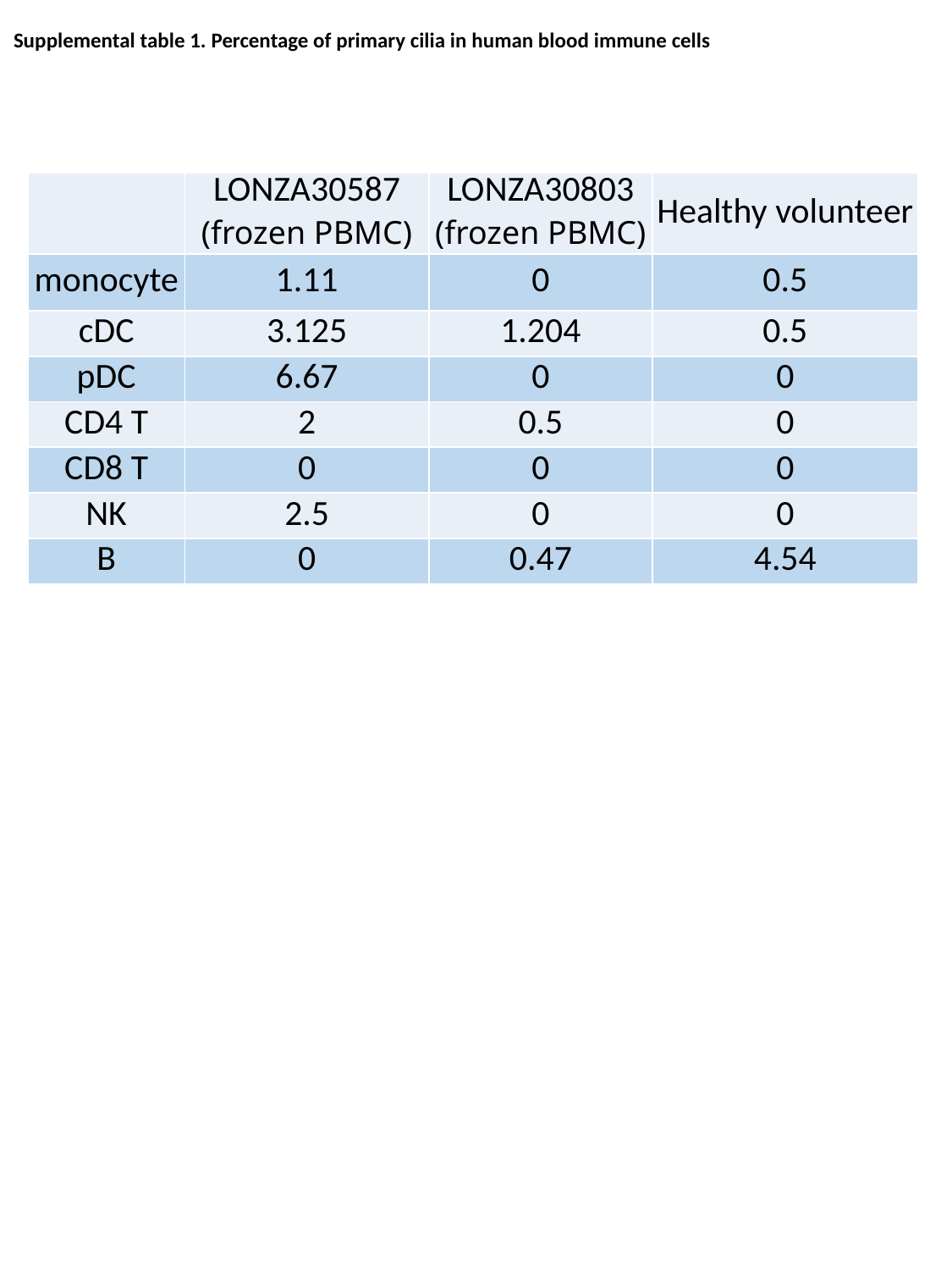

Supplemental table 1. Percentage of primary cilia in human blood immune cells
| | LONZA30587 (frozen PBMC) | LONZA30803 (frozen PBMC) | Healthy volunteer |
| --- | --- | --- | --- |
| monocyte | 1.11 | 0 | 0.5 |
| cDC | 3.125 | 1.204 | 0.5 |
| pDC | 6.67 | 0 | 0 |
| CD4 T | 2 | 0.5 | 0 |
| CD8 T | 0 | 0 | 0 |
| NK | 2.5 | 0 | 0 |
| B | 0 | 0.47 | 4.54 |
